# Kinetic model for the desensitization of G protein-coupled receptor

**DOI:** 10.1101/2024.04.01.587529

**Authors:** Won Kyu Kim, Yoonji Lee, Seogjoo J. Jang, Changbong Hyeon

## Abstract

GPCR desensitization is a general regulatory mechanism adopted by biological organisms against overstimulation of G protein-coupled receptors. Although the details of mechanism aren extensively studied, it is not easy to gain an overarching understanding of the process constituted by a multitude of molecular events with vastly differing time scales. To offer a semi-quantitative yet predictive understanding of the mechanism, we formulate a kinetic model for the G protein signaling and desensitization by considering essential biochemical steps from ligand binding to receptor internalization. The internalization followed by the receptor depletion from the plasma membrane attenuates the downstream signal. Together with the kinetic model, an approximated form of expression derived for the dose-response clarifies the role played by the individual biochemical processes and allows us to identify three distinct regimes for the downregulation that emerge from the balance between phosphorylation, dephosphorylation and the cellular level of *β*-arrestin.

## I. INTRODUCTION

G protein-coupled receptors (GPCRs) are one of many cell surface receptors in living organisms. Their ability to filter numerous extracellular sensory and physiological signals and relay those signals to intra-cellular downstream effector molecules makes GPCRs one of the most important therapeutic targets [1–4].

The agonist binding-induced conformational change in GPCRs enables them to accommodate G proteins and form GPCR/G protein ternary complexes [2]. The ternary complex itself acts as a guanine nucleotide exchange factor (GEF), such that the GEF activity of GPCR/G protein complex promotes the GDP/GTP exchange in G protein [5, 6], which in turn decompose the G protein into G*α* and G*βγ* subunits and releases them into cellular milieu. Multiple subtypes of G*α*, which include G*s*, G*q*, and G*i*, transmit signals of agonist binding by means of second messengers such as cAMP, IP_3_, DAG, and Ca^2+^. Among them, the cellular level of cAMP can be detected using luciferase, enabling to quantify the strength of down-stream signal [7]. Over the past decades, a variety of downstream effector pathways and their connectivity have been elucidated [8], and it has been suggested that the conformational plasticity of GPCRs [9] and GPCR-transducer complexes also contribute to biased signaling and complexities of GPCR signaling [10, 11].

Along with a multitude of signaling pathways, mechanisms to regulate overstimulation of receptors, broadly referred to as GPCR *desensitization* or equivalently known as *drug tolerance* as a longer term consequence, are at work across the entire GPCR super-family. For instance, olfactory receptors display non-monotonic dose-response, such that the intensity of olfactory signal relayed from the receptors drops below its maximum value at high odorant concentrations [12– 14]. A persistent administration of morphine that targets *μ*-opioid receptor downregulates the signal, requiring higher dosage of morphine to elicit the original level of response [15–18]. The desensitization mechanism is activated in response to repeated or prolonged stimulation of GPCR [19–21]. Constitutively active form of GPCR increases the chance of phosphorylation by G protein-coupled receptor kinases (GRKs) or other kinases (cAMP-dependent PKA and lipid signaling-activated PKC), and facilitates the binding of *β*-arrestin to the phosphorylated receptor, blocking G protein interaction with a receptor [22–24]. The receptor internalization and sequestration into endosome [25] result from the *β*-arrestin bound receptor. Either degradation of the internalized receptors without recycling or downregulation of receptor gene transcript reduces receptor density on the cell surface plasma membrane without a change in receptor affinity [15, 25], which then even elicit shrinkage of cell volume [18, 26].

Mechanism of GPCR desensitization has been known and extensively studied over the past decades [20–22, 24, 25, 45–51]; however, many simultaneous kinetic processes with vastly differing time scales make comprehensive assessment of the overall process challenging [52]. In this work, by noting that pre-equilibration of ligand-receptor and ternary complex formation occur faster than the receptor internalization and downregulation, we formulate a minimal kinetic model to offer a semi-quantitative yet predictive understanding of the GPCR desensitization and down-regulation.

## II. MECHANISM OF G PROTEIN SIGNALING AND DESENSITIZATION

In this section, we go through the biochemical steps constituting G-protein signaling and GPCR desensitization depicted in Fig. 1 along with the parameters in Table I, so as to formulate the amplitude of intracellular signal in response to agonist ligand binding.

**TABLE I:** Parameters associated with G protein signaling and desensitization.

| Parameter | Value | Description | Reference |
| --- | --- | --- | --- |
| $G_o$ | $\sim (200 - 3000) \text{ nM}$ | G protein concentration<sup>1</sup> | [27–29] |
| $R_o^i$ | $\sim (1 - 500) \text{ nM}$ | GPCR concentration<sup>2</sup> | [29–35] |
| $K_D$ | $\sim 750 \text{ nM}$ | Binding affinity of GDP-bound G protein | [36] |
| $[\beta A]$ | $\sim 80 \text{ nM}$ | β-arrestin concentration | [34, 35] |
| $K_\beta$ | $\sim (0.1 - 2) \text{ nM}$ | Binding affinity of β-arrestin to phosphorylated GPCR<sup>3</sup> | [37] |
| $V_E (= k_{\text{ex}}[LR^*])$ | $\sim (0.012 - 85) \text{ sec}^{-1}$ | Catalytic rate<sup>4</sup> | [38] |
| $K_E (= K_D)$ | $\sim 750 \text{ nM}$ | Binding affinity of active GPCR to $G_D$<sup>4</sup> | |
| $V_A (= k_{\text{hyd}}\text{RGS}_o)$ | $\sim 0.007 \text{ sec}^{-1}$ | Catalytic rate | [39] |
| $K_A$ | $\sim 30 \text{ nM}, \sim (2 - 600) \text{ nM}$ | Binding affinity of RGS to $G_T$<sup>5</sup> | [40, 41] |
| $V_{\text{max}}^P$ | $t_{1/2} \sim 0.2 \pm 0.05 \text{ min (TRHR)}$<br>$t_{1/2} \sim 1 \pm 0.1 \text{ min } (\beta 2\text{AR})$ | GRK site phosphorylation ($t_{1/2} \sim 1/V_{\text{max}}^P$) | [42] |
| $K_{\text{phos}}$ | $8.3 \pm 1.9 \text{ nM (TRHR)}$<br>$40 \pm 11 \text{ nM } (\beta 2\text{AR})$ | EC50 GRK site phosphorylation | [42] |
| $V_{\text{max}}^{DP}$ | $t_{1/2} \sim 16.2 \pm 1.9 \text{ min (TRHR)}$<br>$t_{1/2} \sim 8.8 \pm 1.9 \text{ min } (\beta 2\text{AR})$ | GRK site dephosphorylation ($t_{1/2} \sim 1/V_{\text{max}}^{DP}$) | [42] |
| $k_{\text{hyd}}$ | $\sim 2.3 \times 10^{-4} \text{ nM}^{-1}\text{sec}^{-1}$ | from $k_{\text{hyd}}\text{RGS}_o \approx 7 \times 10^{-3} \text{ sec}^{-1}$ and $\text{RGS}_o \approx 30 \text{ nM}$ | |
| $k_{\text{ex}}$ | $\sim 1.2 \times 10^{-3} \text{ nM}^{-1}\text{sec}^{-1}$ | from $k_{\text{ex}}[LR^*]_{ss} \approx 0.012 \text{ sec}^{-1}$ and $[LR^*]_{ss} \approx 10 \text{ nM}$ | |
| $\kappa_{\text{AE}}$ | $\sim 0.194$ | $k_{\text{hyd}}/k_{\text{ex}}$ | |
| $K_A^f$ | $\sim 10^{-2}$ | $K_A/G_o \sim 30 \text{ nM}/3000 \text{ nM}$ | |
| $K_E^f$ | $\sim 0.25$ | $K_E/G_o \sim 750 \text{ nM}/3000 \text{ nM}$ | |

**FIG. 1:**
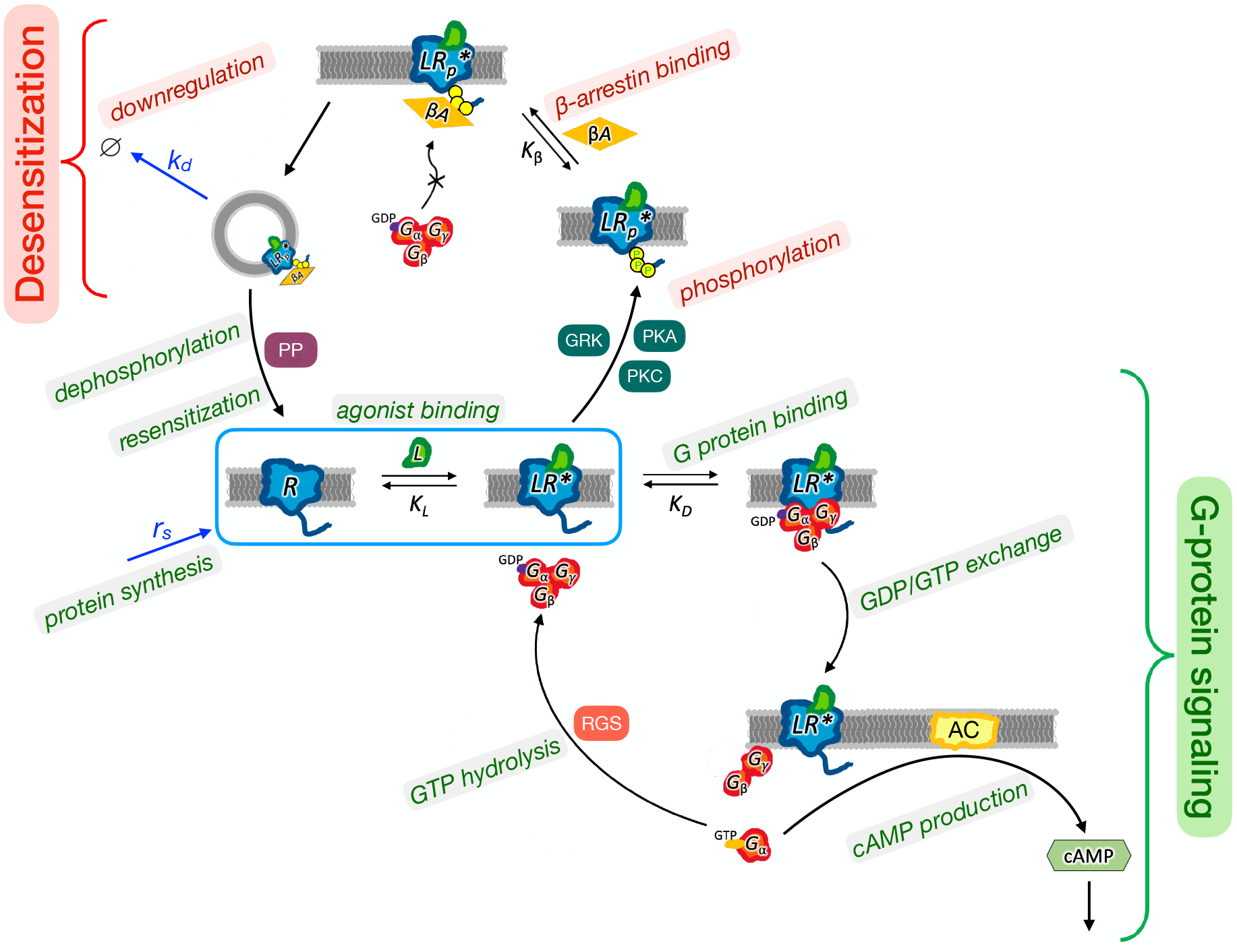
G protein signaling pathway and GPCR desensitization, which includes the G protein signaling that produces cAMP and the desensitization process involved with GPCR phosphorylation, *β*-arrestin binding, internalization and downregulation. Also shown are GEF activity-mediated GDP/GTP exchange, and GTP hydrolysis mediated by GAP activity of RGS, recycling (resensitization) of GPCRs via dephosphorylation, and protein synthesis.

### Receptor activation

G protein signaling starts with the binding of hormone or agonist ligand (*L*) to a receptor on the cell surface (enclosed in the cyan box in Fig. 1)

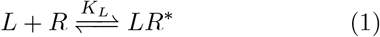

Here, *R* is the inactive form of receptor, and *LR*^*\**^ represents an active conformation of receptor bound with an agonist ligand. The dissociation constant (binding affinity) of this ligand-receptor complex,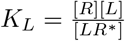 with [*L*] the ligand concentration, is used to relate the concentration of active complex [*LR*^*\**^] with the concentration of free receptor [*R*] and non-dimensionalized ligand concentration *c*_*L*_ *≡* [*L*]*/K*_*L*_ as [*LR*^*\**^] = *c*_*L*_[*R*].

### G protein activation

The active receptor (*LR*^*\**^) can accommodate GDP-bound form of G protein (*G*_*D*_) to form a ternary complex, *LR*^*\**^ *· G*_*D*_, with the binding affinity 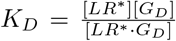. The spontaneous release of GDP from G protein in isolation is extremely slow; yet, the GEF activity of the GPCR/G protein complex can accelerate the exchange of GDP to GTP [5, 6], which leads to the following reaction scheme:

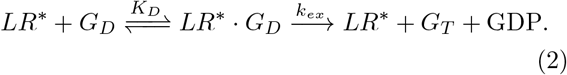

Upon GDP/GTP exchange, the heterotrimeric G protein complex is decomposed into GTP-bound G*α* (*G*_*T*_) and G*βγ* subunits and they are released from the binding cleft of the *LR*^*\**^, stimulating the effectors in the downstream process. The imbalance of *G*_*T*_ and *G*_*D*_ populations (or GTP and GDP) caused by the GEF activity-mediated exchange is restored by regulators of G protein-signaling (RGS) displaying GTPase activating protein (GAP) activity that converts *G*_*T*_ back to *G*_*D*_ via GTP hydrolysis.

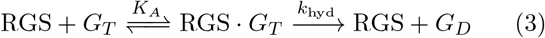

The reaction current associated with the GDP/GTP exchange and the reverse current to balance the cellular concentration of guanosine nucleotides are modeled in a Michaelis-Menten (MM) form:

- 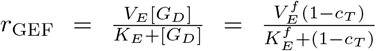 with 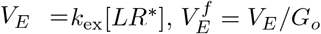 and 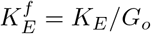.
- 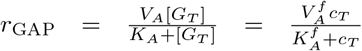 with 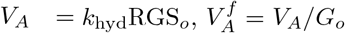 and 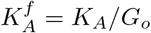.

With the cellular homeostasis in mind, we calculate the cellular concentration of *G*_*T*_ at steady states 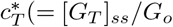 with *G*_*o*_ = [*G*_*T*_] + [*G*_*D*_]) by balancing the two reaction currents, i.e., 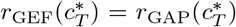 (see Eq. A8 in Appendix A), which yields

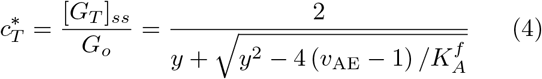

where

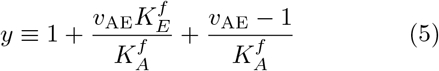

with 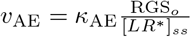 and *κ*_AE_ = *k*_hyd_ */k*_ex_.

### GPCR phosphorylation

When a receptor is subject to prolonged activation, GRKs (and PKA or PKC) phosphorylate serines and threonines in the C-terminal tail or 3rd intracellular loop of the receptor, transforming *LR*^*\**^ into 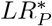 [53]. The kinase-mediated ATP hydrolysis which transfers a phosphate group to the target protein (receptor) is written as

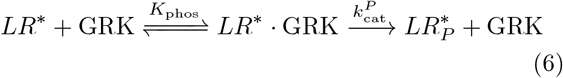

This allows one to model the rate of phosphorylation using MM type kinetics,

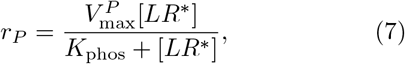

where 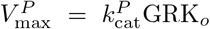 with GRK_*o*_ denoting the cellular level of GRK, and the cellular level of ATP is implicit.

### Desensitization by *β*-arrestin binding and downregulation

Binding of *β*-arrestin to the phosphorylated receptor with its concentration [*βA*]

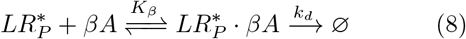

defines the binding constant 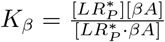. For convenience, we introduce another non-dimensionalized variable *c*_*β*_ *≡* [*βA*]*/K*_*β*_. The ternary complex, 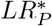, occludes the binding site for G protein (*G*_*D*_) [54, 55]. This engenders a short-term *desensitization* of the receptor or internalization leading to long-term downregulation due to receptor degradation with a rate *k*_*d*_ which ranges from *hours* to *days* [20].

While receptor depletion from the plasma membrane occurs due to the *β*-arresitin binding-induced endocytosis [26], it is reasonable to assume that the receptors are still synthesized and replenished with a rate *r*_*s*_, which leads to the following evolution equation for the receptor density on the cell surface:

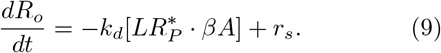

### Dephosphorylation and resensitization of receptor

In the longer time scale, it was suggested that typically only after the receptor internalization, protein phosphotases (PPs) are at work in an acidic endosomal compartment to dephosphorylate, resensitize and restore the receptor back to the cell surface [48, 50, 56, 57]. Thus, the reaction scheme for the dephosphorylation of the ternary complex of 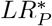 mediated by PPs is expressed as

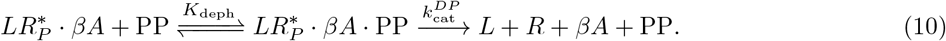

Similarly to the rate of phosphorylation (Eq. 7), the rate of dephosphorylation is modeled as

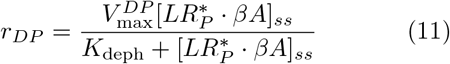

with 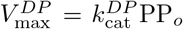 and PP_*o*_ denoting the cellular level of phosphatase.

When the reaction cycle associated with the desensitization is at steady state, the current flowing in the cycle should be constant, balancing the rates between the phosphorylation and dephosphorylation, i.e., *r*_*P*_ *≈ r*_*DP*_. Thus,

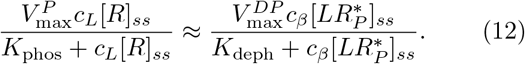

This enables to express 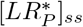 in terms of [*R*]_*ss*_:

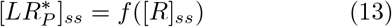

where

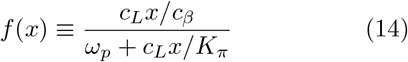

with

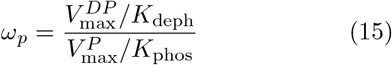

and

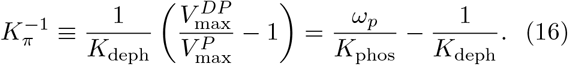

The *ω*_*p*_, the ratio of Michaelis rates of dephosphorylation and phosphorylation defined in Eq. 15, will be shown to play a key role in determining the dose-response.

### Mass balance

In the absence of receptor depletion, more precisely in the limit of time scale separation between the receptor depletion and the dynamics of ligand-induced G protein signaling, the total concentration (number) of receptors on the plasma membrane should satisfy the mass balance:

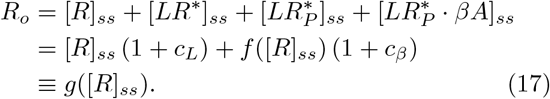

Thus, it follows from the mass balance that [*R*]_*ss*_ can formally be expressed in terms of *R*_*o*_

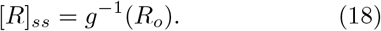

The expressions of the other receptor states are also obtained by employing Eqs. 14 and 18.

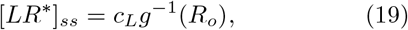

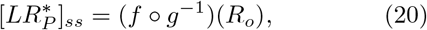

and

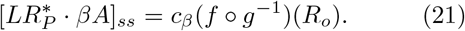

### G protein signaling

The 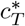 is directly linked to the activation of adenylyl cyclase (AC) that leads to the production of cAMPs and downstream signal,

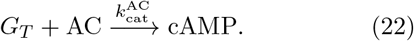

From an assumption that the G protein signaling (*δS*) above the basal level is proportional to the cellular level of cAMP [14, 58, 59], the intensity of signaling is given by

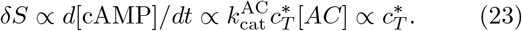

More detailed expression of *δS* for further quantitative analysis is derived and discussed below.

## III. RESULTS AND DISCUSSION

The amplitude of G protein signaling measured after an incubation time *t, δS*(*t*), would be proportional to the cAMP being produced (Eq. 22). From Eqs. 4 and 23, the amplitude of signaling reads

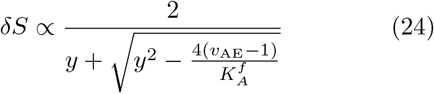

where

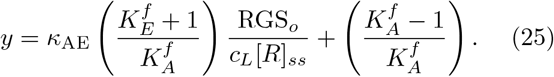

[*R*]_*ss*_ can be expressed in terms of *R*_*o*_ by rewriting Eq. 17,

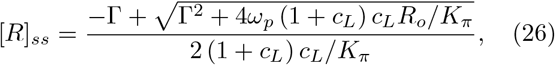

with

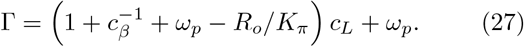

Incidentally, the receptor concentration *R*_*o*_ obeys Eq. 9 and it changes slowly from its initial concentration to the steady state value due to the downregulation and synthesis (or resensitization). Eq. 21 allows us to cast Eq. 9 into a form of ordinary differential equation,

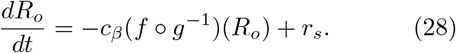

Although the explicit expression of (*f ○ g*^*−*1^)(*R*_*o*_) is lengthy and complicated, it is straightforward to solve Eq. 28 numerically. The numerical solution of *R*_*o*_(*t*) acquired for a given set of parameters (*ω*_*p*_, *c*_*L*_, *c*_*β*_) is plugged into Eqs. 26, 27, which eventually offers the full numerics of *δS* in Eq. 24 and the plots in Fig. 2.

**FIG. 2:**
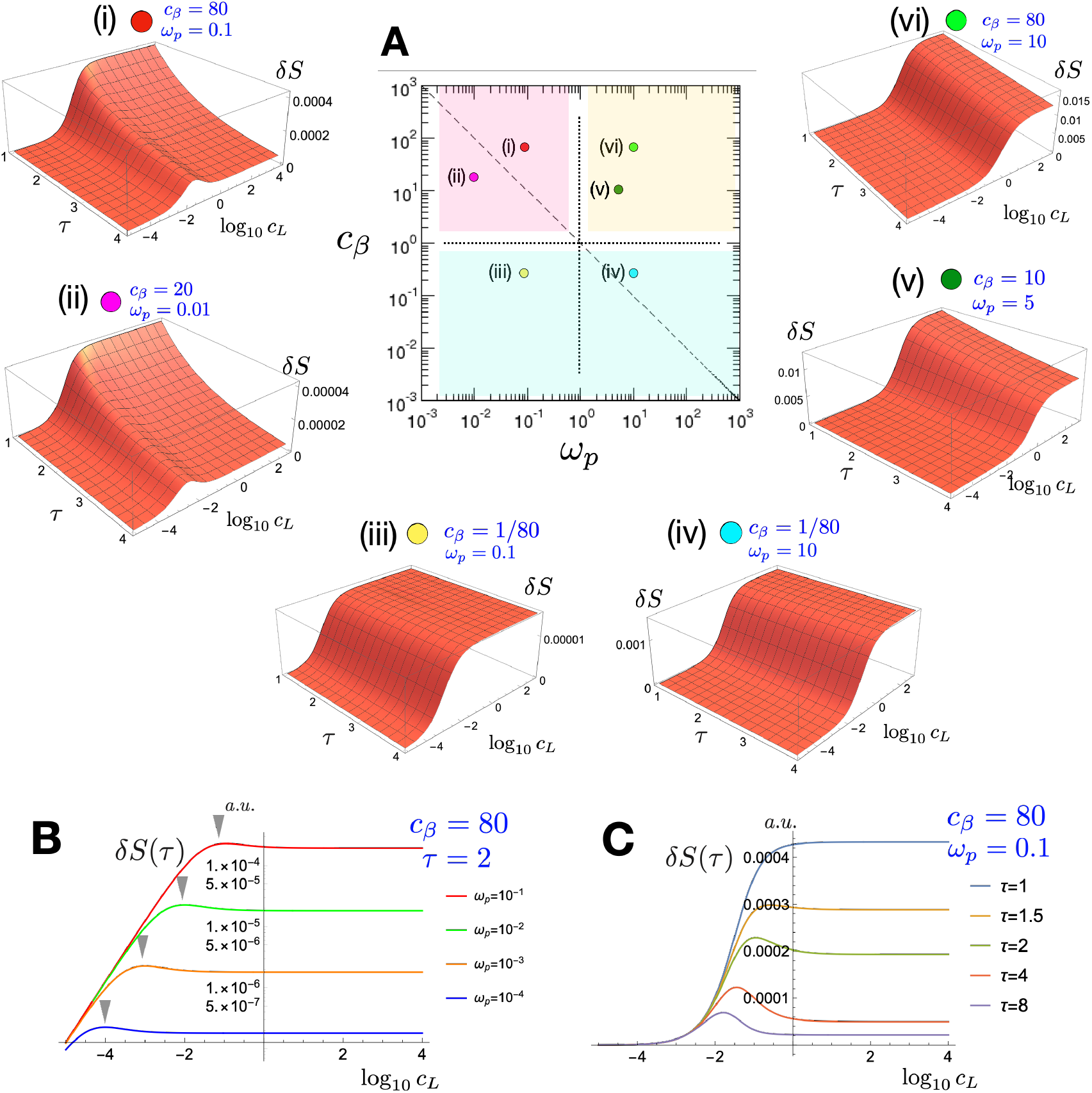
Variation of *δS* with *c*_*L*_ and measurement (or incubation) time (*τ* = *k*_*d*_*t*) calculated using Eqs. 32. (A) *δS*(*c*_*L*_; *τ*) calculated for (i) *c*_*β*_ = 80, *ω*_*p*_ = 0.1, (ii) *c*_*β*_ = 20, *ω*_*p*_ = 0.01, (iii) *c*_*β*_ = 1*/*80, *ω*_*p*_ = 0.1, (iv) *c*_*β*_ = 1*/*80, *ω*_*p*_ = 10. (v) *c*_*β*_ = 10, *ω*_*p*_ = 5. (vi) *c*_*β*_ = 80, *ω*_*p*_ = 10. The panel in the middle depicts the sets of parameters (*ω*_*p*_, *c*_*β*_) used for the calculations. The parameter space giving rise to three distinct kinetic behaviors of downregulation (the cases (*a*), (*b*), and (*c*) discussed in Eq. 38) are demarcated with different colors. (B) *δS*(*τ* = 2) versus *c*_*L*_ for varying *ω*_*p*_ with *c*_*β*_ = 80. The position of *δS*_max_ for varying *ω*_*p*_ value is highlighted with inverted triangles. Note that *δS* is plotted in logarithmic scale. (C) *δS*(*τ*) versus *c*_*L*_ for varying *τ* with *ω*_*p*_ = 0.1 and *c*_*β*_ = 80. The parameters used for the plots are 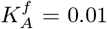, *K*_phos_ = 40 nM, *K*_deph_ = 40 nM, *K*_*D*_ = 750 nM, RGS_*o*_ = 100 nM, 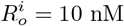, and *r*_*s*_ = *k*_*d*_*/*6 (see Table I).

Equation 24 along with Eqs. 25–27 is the key expression of this study for the amplitude of G protein signaling without any approximation. Although plotting *δS* numerically as a function of *c*_*L*_ and *ω*_*p*_ (see Fig. 2) is straightforward, the complexity of the expression prevents one from gaining physical insights of the G protein signaling and desensitization. To this end, we approximate *δS* (Eq. 24) and derive a physically more interpretable expression, *δS*^*†*^ (Eq. 31).

First, using the condition of 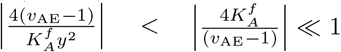 for *v*_*AE*_*≪* 1 and 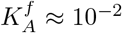 (Table I), we obtain

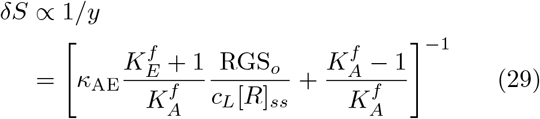

Second, a condition of 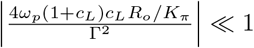, which is valid for *c*_*L*_ *≪* 1, further approximates Eq. 26 as

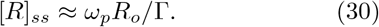

This simplifies *δS* to

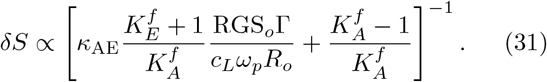

By rewriting Eq. 31 along with Eq. 27, we obtain *δS*^*†*^(*c*_*L*_; *t*), a physically interpretable form of MM-like dose-response behavior weighted with a factor 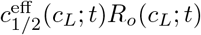

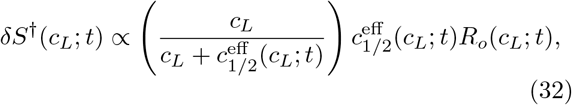

where the middle point of signaling, corresponding to EC_50_ value, is given by

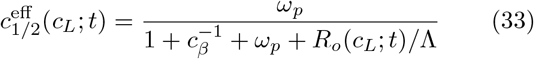

with

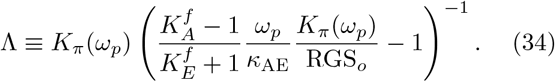

The amplitude and EC_50_ value of *δS*^*†*^ are adjusted by the dose- and incubation time-dependent receptor population *R*_*o*_(*c*_*L*_; *t*). The evolution equation of *R*_*o*_ (Eq. 9) can be approximated as

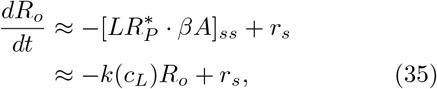

which yields

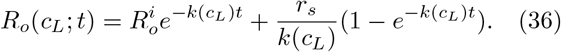

Here,

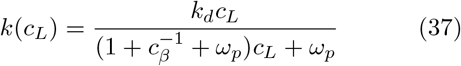

is acquired from the relations 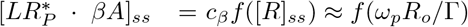 and Eq. 14.

The MM-like structure of *δS*^*†*^ (Eqs. 32) along with 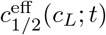 (Eq. 33), *R*_*o*_(*c*_*L*_; *t*) (Eq. 36), and *k*(*c*_*L*_) (Eq. 37) allows us to understand the behaviors of *δS* (Eq. 24) plotted in Fig. 2 for varying sets of parameters and to dissect the contribution of the individual kinetic process to the downstream signal and desensitization. The expressions of 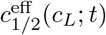 (Eq. 33) and *k*(*c*_*L*_) (Eq. 37) suggest that *ω*_*p*_ (Eq. 15), which dictates the balance between dephosphorylation and phosphorylation, and *c*_*β*_, the cellular level of *β*-arrestin, are the two key parameters that determine the dose-response behavior. The effects of the parameters associated with other processes, such as GEF/GAP activity-mediated processes (*v*_AE_, *K*_*A*_, and *K*_*E*_), binding affinity of *G*_*D*_ to a receptor (*K*_*D*_) are consolidated into a single term Λ (Eq. 34) along with the parameters involving phosphorylation/dephosphorylation, i.e., *ω*_*p*_ (Eq. 15), *K*_*π*_ (Eq. 16). For *ω*_*p*_ *<* 1, phosphorylation occurs more frequently than dephosphorylation and the chance of desensitization increases. Conversely, *ω*_*p*_ *>* 1 would lead to resensitization. Although *β*-arrestins are replete in the cellular milieu (*c*_*β*_ *≈* (40 *−* 800) *≫* 1) [34, 35] (see Table I), we expand *c*_*β*_ to a broader range, 10^*−*5^ *< c*_*β*_ *<* 10^4^ to elucidate the role played by *β*-arrestin on G protein signaling and desensitization (Fig. 2).

The depletion rate of receptors from the plasma membrane, *k*(*c*_*L*_), for the limit of *c*_*L*_ *≫* 1 can be approximated to three cases of (*a*), (*b*), and (*c*), depending on the values of *ω*_*p*_ and *c*_*β*_.

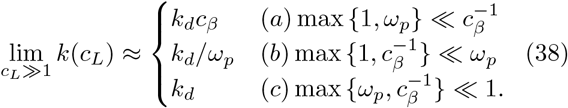

The case (c) corresponds to the situation that phosphorylation is dominant (*ω*_*p*_ *≪* 1) and cellular level of *β*-arrestin is high (*c*_*β*_ *≫* 1), which gives rise to the fastest decay rate (*k*_*d*_). In other cases ((*a*) and (*b*)), the downregulation is suppressed.

The Figure 2A shows *δS*(*c*_*L*_, *τ*) calculated under 6 different sets of parameters: (i) and (ii) calculated with *c*_*β*_ *≫* 1, *ω*_*p*_ *≪* 1, corresponding to the case (*c*) in Eq. 38, display downregulations with a rate *k*_*d*_, and non-monotonic variations of dose-response; (iii) and (iv) with *c*_*β*_ *≪* 1 belong to the case (*a*) where the decay rate is reduced by 80 fold, i.e., *k*_*d*_ *→ k*_*d*_*c*_*β*_ = *k*_*d*_*/*80; (v) and (vi) with *ω*_*p*_ *>* 1 belong to the case (*b*) with the decay rate being reduced to *k*_*d*_ *→ k*_*d*_*/ω*_*p*_. The decay rates for (v) and (vi) are decreased to *k*_*d*_*/*5 and *k*_*d*_*/*10, respectively.

Of particular interest is the peculiar shapes of dose-response obtained for the case (*c*) (*c*_*β*_ *≫* 1 and *ω*_*p*_ *≪* 1). If the incubation times are comparable to the decay time, namely, *τ* (= *k*_*d*_*t*) *∼ 𝒪* (1), *R*_*o*_(*c*_*L*_; *t*) decreases linearly with *c*_*L*_ for *c*_*L*_ *≪ ω*_*p*_ and becomes *c*_*L*_-independent for *c*_*L*_ *≫ ω*_*p*_ (Appendix B). The hyperbolic saturation of the term inside the parentheses in Eq. 32 and the decrease of *R*_*o*_(*c*_*L*_; *t*) with *c*_*L*_ yield a non-monotonic variation of *δS*(*τ*) peaked around at *c*_*L*_ *∼ ω*_*p*_, as exemplified by *δS*(*τ* = 2) plotted for various *ω*_*p*_(*≪* 1) with *c*_*β*_ = 80 in Fig. 2B. In fact, this type of non-monotonic response to ligand concentration after a finite incubation time is seen in the literature (see Fig. 3). Next, it is noteworthy that the overall amplitude of *δS* is decided by *ω*_*p*_. The smaller *ω*_*p*_ leads to a smaller *δS*. In fact, this is consistent with the notion that the downstream signaling is suppressed under phosphorylation dominant condition (*ω*_*p*_ *≪* 1).

**FIG. 3:**
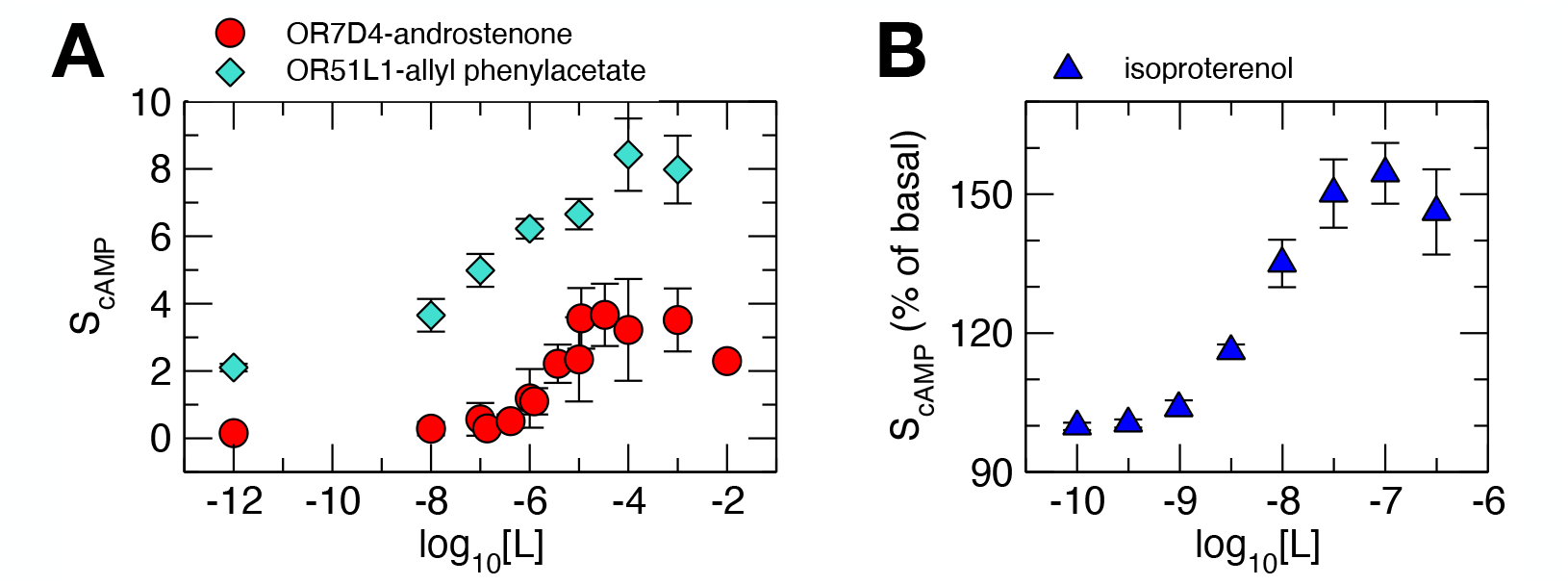
Examples of *non-monotonic* response of G protein signaling to the increasing concentration of ligand. (A) Dose-response of olfactory receptor neuron that expresses only a single type of olfactory receptor (OR). The level of cAMP stimulation was measured using luciferase against an increasing concentration of cognate odorants. Plotted are two pairs of OR versus cognate odorant data taken from Ref. [13]: OR7D4 versus androstenone (red circle), and OR51L1 versus allyl phenylacetate (cyan diamond). (B) Dose-response curves for cAMP accumulation in sympathetic neurons measured after 10 min incubation time with an increasing concentration of isoproterenol (non-selective *β*-adrenergic receptor agonist used for the heart related treatment) [60].

Finally, two scenarios giving rise to a system effectively free from downregulation could further illuminate the findings of this work. First, in the dephosphorylation dominant regime (*ω*_*p*_ *>* 1), the chance of resentization increases and *k*(*c*_*L*_) *≈* (*k*_*d*_*/ω*_*p*_)*c*_*L*_*/*(*c*_*L*_+1) saturates to *k*_*d*_*/ω*_*p*_ for *c*_*L*_ *≫* 1 (case (*b*) in Eq.38). As demonstrated in Fig. 2A-(v) and (vi) obtained with *ω*_*p*_ *>* 1 and *c*_*β*_ *>* 1, *δS*(*τ*)s are no longer non-monotonic with *c*_*L*_. Next, if *β*-arrestin level is low (*c*_*β*_ *≪* 1), then the precursor complex of the internalization and degradation, i.e., 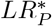 can no longer be formed, and the EC_50_ value is effectively written as 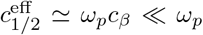 Consequently, *δS*(*c*_*L*_, *τ*) effectively devoid of downregulation is obtained as depicted in Fig. 2A-(iii) and (iv). Taken together, our kinetic model clarifies that the receptor phosphorylation and high cellular level of *β*-arrestins are the two key requirements for the GPCR desensitization [20].

## IV CONCLUDING REMARK

While the reaction scheme represented by Fig. 1 capture the major mechanism of GPCR signaling and desensitization, it is known that the actual cellular processes are even more complicated. Indeed, the detailed mechanism and regulatory pathways involving receptor phosphorylation [61, 62], *β*-arrestin binding [63, 64], and internalization [20, 65], resensitization and recycling through dephosphorylation [20, 66, 67] may not only be agonist-selective [67–69], but also depends on receptor subfamily, cell and tissue types [20, 70, 71]. Specifically, two different agonists could stabilize a receptor into two distinct conformations, triggering different phosphorylation pathways [69]. Despite a long history of study on the subject, systems level insight into GPCR desensitization is difficult to gain, as intracellular signal transductions in general are featured with substantial complexity. Even if one has full knowledge of the associated reaction network along with the binding constants and rate constants, it is not so easy to grasp how and what type of cell physiology emerges from the network and which biochemical steps are the key factors responsible for the resulting process [52, 72].

In this study, confining ourselves to a prototypical mechanism of G protein signaling and homologous desensitization [22, 25] (Fig. 1), we have derived an expression of the incubation time-dependent dose-response of downstream signaling. The dose-response behaviors of G protein signaling explored in this study (Fig. 2) and the MM-like expression derived from our formulation clarify that phosphorylation and *β*-arrestin binding are the two main processes responsible for shaping a drug tolerance (a suppressed downstream signaling at high agonist concentration), a dose-response widely observed in biochemical assays (Fig. 3). Although we demonstrated our theoretical consideration on GPCR desensitization by limiting to a prototypical desensitization pathway, it is straight-forward to extend our formulation to include other pathways such as phosphorylation by cAMP-induced activation of PKA [73, 74], which requires including a feedback regulation, and dephosphorylation without internalization [56, 75] and so forth.

To recapitulate, our theoretical study on GPCR desensitization not only offers a comprehensive systems-level understanding of the interplay among a series of molecular events, but also highlights the importance of the balance in phosphorylation/dephosphorylation cycle and the cellular level of *β*-arrestin in regulating the overstimulation of G protein signaling.

## Acknowledgments

This study was supported by KIAS Individual Grants CG076002 (W.K.K.), and CG035003 (C.H.), and by the US National Science Foundation (CHE-1900170 to S.J.J.). We thank the Center for Advanced Computation in KIAS for providing the computing resources.

## Appendix A: Goldbeter–Koshland zero-th order ultrasensitivity and derivation of 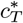 in Eq. 4

The key expression of this study, 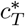 (Eq. 4), is derived from the balance between the GEF and GAP activity-mediated processes. The expression bears the same mathematical structure with that of the phosphorylation-dephosphorylation cycle, formulated by Goldbeter and Koshland, which is known to yield ultrasensitive responses of substrates to the relative amount of two opposing enzymes [76, 77].

Here we derive a general expression equivalent to 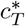 by considering the chemical state of a receptor catalyzed by phosphatase (*P*) with concentration *P*_*o*_ and kinase (*K*) with concentration *K*_*o*_. When a large number of substrates (receptors) are present, such that the total concentration of phosphorylated (*Z*_*p*_) and un-phosphorylated receptors (*Z*) is greater than *K*_*o*_ and *P*_*o*_, i.e., *Z*_*tot*_ *≫ K*_*o*_, *P*_*o*_ with *Z*_*tot*_ = [*Z*] + [*Z*_*p*_], we can assume that the system is in the Michaelis-Menten (MM) regime. The interconversion of the receptor between *Z*_*p*_ and *Z* is written as

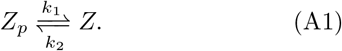

*Z*_*p*_ is dephosphorylated with the rate

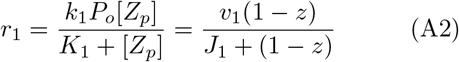

where *z* = [*Z*]*/Z*_*tot*_ with *v*_1_ = *k*_1_*P*_*o*_, and *J*_1_ = *K*_1_*/Z*_*tot*_. *Z* is phosphorylated with the rate

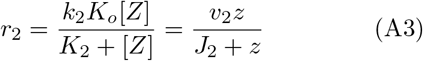

where *v*_2_ = *k*_2_*K*_*o*_ and *J*_2_ = *K*_2_*/Z*_*tot*_. The concentration of unphosphorylated receptor at steady state, *z*^*\**^ = [*Z*]_*ss*_*/Z*_*tot*_, is decided from *r*_1_(*z*^*\**^) = *r*_2_(*z*^*\**^),

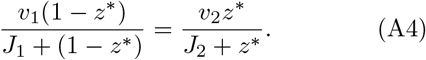

The state of the receptor changes in the range of 0 *≤ z*^*\**^ *≤* 1 in response to the relative rate of de-phosphorylation and phosphorylation (or the relative amount of kinase and phosphatase) in the cell (*v*_1_*/v*_2_ = *k*_1_*P*_*o*_*/k*_2_*K*_*o*_). Physically, it is expected that the receptors are in the unphosphorylated state when the rate of dephosphorylation is greater than that of phosphorylation, and vice versa. This switch-like transition of *z*^*\**^ between *z*^*\**^ *≈* 0 and *z*^*\**^ *≈* 1 is further clarified by rewriting Eq. A4 in the following form with an assumption of *J*_1_ = *J*_2_ = *J*,

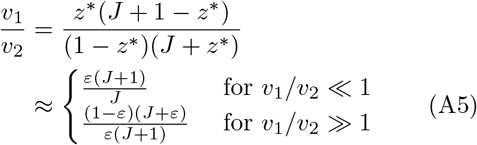

**FIG. A1:**
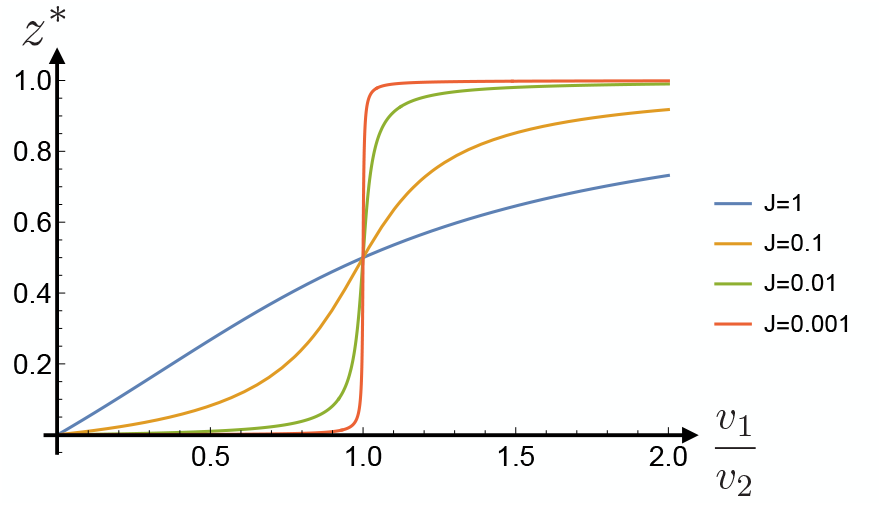
Plots of *z*^*\**^ versus *v*_1_*/v*_2_ for varying *J* (Eq. A5). This demonstrates the ultrasensitive response of substrate to the two opposing enzymes, especially at small *J* .

with *ε ≪* 1, and can more explicitly be demonstrated by plotting *z*^*\**^ as a function of *v*_1_*/v*_2_ (Fig. A1). The first and the second derivatives at the transition point *v*_1_*/v*_2_ = 1

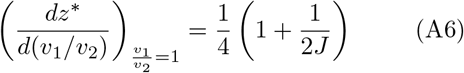

and

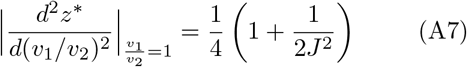

indicate that small *J* value sharpens the transition between *z*^*\**^ = 0 and *z*^*\**^ = 1.

Solving Eq. A4 for *z*^*\**^ yields the Goldbeter’s formula for the zero-th order ultrasensitivity:

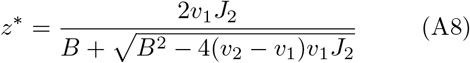

where *B* = *v*_1_*J*_2_ + *v*_2_*J*_1_ + *v*_2_ *− v*_1_. Note that only one of the two solutions from the quadratic equation is physically relevant since 0 *≤ z*^*\**^ *≤* 1. The term inside the square root of Eq. A8

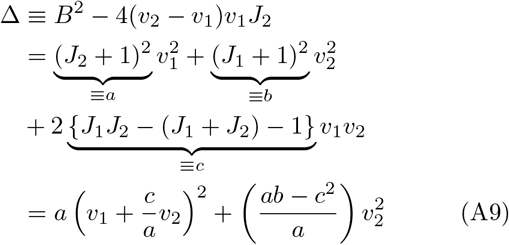

is positive for all values of *v*_1_, *v*_2_, *J*_1_, *J*_2_ *>* 0, because *a, b >* 0 and *ab − c*^2^ = 4*J*_1_*J*_2_(*J*_1_ + *J*_2_ + 1) *>* 0.

Along with the inequalities 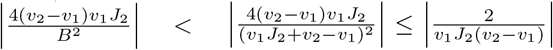, if the system is either in the dephosphorylation or phosphorylation dominant regime 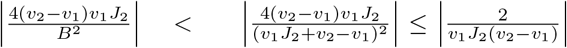 is guaranteed, which simplifies the expression of *z*^*\**^ to

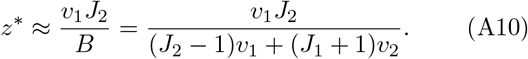

## Appendix B: Receptor population, *R*_*o*_(*c*_*L*_; *τ*)

For *c*_*β*_ *≫* 1 and *ω*_*p*_ *≪* 1, *R*_*o*_(*c*_*L*_; *τ*) with *k*(*c*_*L*_) *≈ k*_*d*_*c*_*L*_*/*(*c*_*L*_ + *ω*_*p*_) (Eq. 37) is approximated at short and long time limits as follows:

1. For short incubation time, *τ* (= *k*_*d*_*t*) *∼ 𝒪*(1),

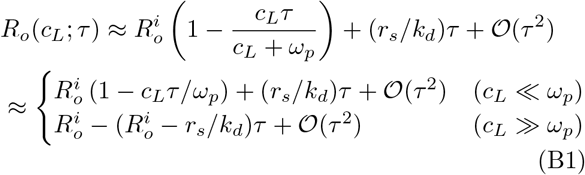
2. For long incubation time, *τ ≫* 1,

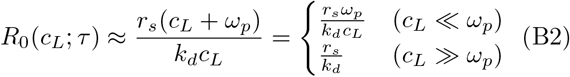

